# In BRCA1 and BRCA2 breast cancers, chromosome breaks occur near herpes tumor virus sequences

**DOI:** 10.1101/2021.04.19.440499

**Authors:** Bernard Friedenson

**Affiliations:** Dept. of Biochemistry and Molecular Genetics, College of Medicine, University of Illinois Chicago

**Keywords:** Breast cancer infection, breast cancer immunity, breast cancer virus, genome biochemistry, nasopharyngeal cancer, herpes viruses, hereditary breast cancer, BRCA1, BRCA2, Burkitt’s lymphoma, homologous recombination, DNA repair, viral cancer, chromosome breaks

## Abstract

Inherited mutations in BRCA1 and BRCA2 genes increase risks for breast, ovarian, and other cancers. Both genes encode proteins for accurately repairing chromosome breaks. If mutations inactivate this function, broken chromosome fragments get lost or reattach indiscriminately. These mistakes are characteristic of hereditary breast cancer. We tested the hypothesis that mistakes in reattaching broken chromosomes preferentially occur near viral sequences on human chromosomes. We tested millions of DNA bases around breast cancer breakpoints for similarities to all known viral DNA. DNA around breakpoints often closely matched the Epstein-Barr virus (EBV) tumor variants HKHD40 and HKNPC60. Almost all breakpoints were near EBV anchor sites, EBV tumor variant homologies, and EBV-associated regulatory marks. On chromosome 2, EBV binding sites accounted for 90% of breakpoints (p<0.0001). On chromosome 4, 51/52 inter-chromosomal breakpoints were close to EBV variant sequences. Five viral anchor sites at critical genes were near breast cancer breakpoints. Twenty-five breast cancer breakpoints were within 1.25% of breakpoints in model EBV cancers. EBV-like sequence patterns around breast cancer breakpoints resemble gene fusion breakpoints in model EBV cancers. All BRCA1 and BRCA2 breast cancers had mutated genes essential for immune responses. Because of this immune compromise, herpes viruses can attach and produce nucleases that break chromosomes. Alternatively, anchored viruses can retard break repairs, whatever the causes. The results imply proactive treatment and prevention of herpes viral infections may benefit BRCA mutation carriers.

## Introduction

Inherited mutations in BRCA1 and BRCA2 genes increase risks for breast, ovarian and other cancers. As one of their many functions, the two genes encode proteins needed to restore broken DNA by homologous recombination. If mutations inactivate either the BRCA1 or BRCA2 gene, then the broken DNA can become pathogenic. Pieces of DNA get lost or reattach at the wrong positions on the original or different chromosomes. In carriers of BRCA1 or BRCA2 gene mutations, these errors lead to chromosome rearrangements and shifts typical of hereditary breast cancers. Landmark studies of Nik-Zainal and colleagues report hundreds of DNA rearrangements and characteristic rearrangement signatures[1-4]. Chromosome rearrangements may be critical events leading to hereditary breast cancers, but we know very little about what causes these events.

About 8% of human DNA probably originated from retroviruses [5] and both endogenous and exogenous retrovirus sequence insertions are a significant source of genome instability. Exogenous human herpesvirus 4 (EBV) can destabilize the genome by inducing DNA breaks. EBV infects over 90% of all adults worldwide. Most hosts remain healthy because their immune systems control the infection, but EBV stays for life as a latent circular episome in host memory B-cells.

EBV associates with a diverse group of malignancies. EBV infection is always present in undifferentiated nasopharyngeal cancer (NPC) [6-8]. Downregulation or mutations in DNA repair genes related to BRCA1-BRCA2 mediated repair pathways are common in NPC [9,10]. Because of repair defects, NPC shows chromosomal shifts and rearrangements, like those seen in hereditary breast cancers.

HKNPC60 and HKHD40 are two EBV tumor virus variants 100% associated with NPC [11]. NPC in human epithelial cells arises after viral DNases induce DNA strand breaks followed by inappropriate gene fusions, such as YAP1-MAML2, PTPLB-RSRC1, and SP3-PTK2 [12]. EBV produces an exonuclease (BGLF5) and an endonuclease (BALF3). These viral products cause DNA strand breaks and genome instability [13,14], characteristic of cancers in EBV infected lymphoid cells [15]. Fortunately, a signaling pathway dependent on BRCA1 and BRCA2 genes prevents EBV from causing malignant transformation [16].

The same EBV-sensitive, BRCA-related pathway is also essential to prevent hereditary breast cancers, suggesting a role for EBV infection in hereditary breast cancers. Some previous literature supports this relationship. EBV infection of breast epithelial cell models facilitates malignant transformation and tumor formation [17]. Epidemiological associations between breast cancer and EBV infection exist in different geographical locations [18,19]. Breast cancer cells from biopsies express gene products from latent EBV infection (LMP-1, −2, EBNA-, and EBER) [20] even after excluding the possibility that the virus comes from lymphocytes [21]. The EBV lytic form in breast cancer associates with a worse outcome [22]. Even though breast cancer malignancies may not have active EBV infection, the virus may still leave its footprints. Carriers of hereditary mutations in BRCA1 and BRCA2 genes would be especially susceptible to damage from viral infection.

Immune deficits in BRCA1 and BRCA2 mutation carriers [23] may allow the reactivation of latent EBV infections or new herpes viral infections. DNA breaks induced by EBV nucleases are then challenging to repair correctly and may become pathogenic. The availability of breast cancer genomic sequences permits testing the possibility that EBV contributes to the incorrect reattachment of broken chromosomes in hereditary breast cancer.

## Materials and Methods

### Breast cancer genomic sequences

#### Characteristics of breast cancers compared to viral cancers

Breast cancer genome sequences were from the COSMIC database curated from an original publication [3,4]. 15/25 breast cancers were stage III, four were stage II, and three had no data. Six breast cancers had typed germ-line BRCA1 mutations, and nineteen had typed germ-line BRCA2 mutations. Blood was the source of normal genes for comparison [4]. All 25 cancers were female ductal breast cancers.

The 25 ductal breast cancers contained many DNA missense mutations, with an average of 1.55 mutations per gene analyzed (range 1.0-2.75). A total of 275,730 mutations had a mean value of 11,029 per cancer, ranging from 1-2.75 per gene examined. The 4316 DNA breakpoints varied from 33 to 396 per different individual cancer with a mean of 173. For all 25 breast cancers, chromosomes 1 and 2 were the most frequent sites of intra-chromosome rearrangements, but the distributions of DNA breakpoints among the various chromosomes were markedly different (Table 1).

**Table 1.** BRCA1 and BRCA2 breast cancers studied

| Sample | BRCA1 /<br>BRCA2<br>mutation<br>status | Analyzed<br>genes | Muta-<br>tions | Breaks | Muta-<br>tions<br>per<br>gene | Muta-<br>tions per<br>break | Chro-<br>some<br>with most<br>breaks<br>"From" | Chro-<br>some<br>with most<br>breaks "To" | Most often<br>intra-chro-<br>mosomal<br>breaks |
| --- | --- | --- | --- | --- | --- | --- | --- | --- | --- |
| PD3890 | BRCA1 | 5978 | 13401 | 269 | 2.24 | 49.8 | 3=4,1 | 11, 16,21 | 1 |
| PD3904 | BRCA1 | 6337 | 13559 | 247 | 2.14 | 54.9 | 6 | 17,18 | 8 |
| PD3945 | BRCA1 | 9203 | 23132 | 114 | 2.51 | 202.9 | 1 | X | 2 |
| PD4005 | BRCA1 | 6943 | 15093 | 185 | 2.17 | 81.6 | 3 | 5 | 1 |
| PD4006 | BRCA1<br>R1835X | 7433 | 20453 | 396 | 2.75 | 51.6 | 1 (only 1) | 22,16 | 2 (387 intra) |
| PD4115 | BRCA1* | 9069 | 22662 | 180 | 2.50 | 125.9 | 2 | 17 | 8 |
| PD4116 | BRCA2 | 7831 | 19958 | 359 | 2.55 | 55.6 | 11 | 17 | 3 |
| PD4836 | BRCA2 | 4782 | 5381 | 80 | 1.13 | 67.3 | 10,11,12 | 18,20 | 10 |
| PD4872 | BRCA2 | 5095 | 5906 | 90 | 1.16 | 65.6 | 2 | 18 | 13 |
| PD4874 | BRCA2 | 8201 | 10668 | 226 | 1.30 | 47.2 | 6 | 20 | 10 |
| PD4875 | BRCA2 | 6262 | 8056 | 140 | 1.29 | 57.5 | 5 | 19 | 7 |
| PD4876 | BRCA2 | 5718 | 8317 | 95 | 1.45 | 87.5 | 5 | 9 | 12 |
| PD4951 | BRCA2 | 3178 | 3283 | 33 | 1.03 | 99.5 | 6 | 8 | 4 |
| PD4952 | BRCA2 | 10806 | 15155 | 332 | 1.40 | 45.6 | 3 | 14 | 14 |
| PD4953 | BRCA2 | 6690 | 8090 | 143 | 1.21 | 56.6 | 4 | 18 | 18 |
| PD4954 | BRCA2 | 5904 | 6605 | 126 | 1.12 | 52.4 | 1 | 8 | 2 |
| PD4955 | BRCA2 | 6654 | 7920 | 100 | 1.19 | 79.2 | 12 | 15 | 1 |
| PD4956 | BRCA2 | 11036 | 14860 | 238 | 1.35 | 62.4 | 7 | 10 | 3 |
| PD4957 | BRCA2 | 3521 | 3520 | 74 | 1.00 | 47.6 | 7=10 | 17=11 | 8 |
| PD4958 | BRCA2 | 8244 | 10956 | 102 | 1.33 | 107.4 | 1 | 1 | 4 |
| PD6406 | BRCA2 | 4976 | 5485 | 191 | 1.10 | 28.7 | 2,4 | 6,8 | 2 |
| PD6416 | BRCA2 | 3364 | 3609 | 71 | 1.07 | 50.8 | 1 | 20 | 7 |
| PD7217 | BRCA2 | 8841 | 11215 | 124 | 1.27 | 90.4 | 3 | 18 | 3 |
| PD8621 | BRCA2 | 7964 | 9617 | 275 | 1.21 | 35.0 | 1 | X | 1 |
| PD8969 | BRCA2 | 7335 | 8829 | 126 | 1.20 | 70.1 | 1 | 8 | 9 |

\*Mutation of uncertain significance

The genes analyzed in BRCA1-associated breast cancers (Table 1) were on average almost twice as likely to be mutated as the genes analyzed in BRCA2 associated breast cancers (p<0.0001). Moreover, an unpaired t-test assuming equal variances found that the means were different (BRCA1=94.5, BRCA2 63.5, p=0.03

In some cases, we also used breast cancer genome data from presumptive mutation carriers. These women were all under age 36 but had not been tested for BRCA mutations.

#### DNA sequence data

We used the NCBI BLAST program (MegaBLAST) and database [24,25] to compare DNA sequences around breakpoints in BRCA1-and BRCA2-mutation-positive breast cancers to all available viral DNA sequences. Virus DNA was from BLAST searches using “viruses (taxid:10239)” with homo sapiens excluded. Gene break-points for inter-chromosomal and intra-chromosomal translocations were obtained from the COSMIC catalog of somatic mutations as curated from original publications [4] and converted to the GrCH38 human genome version. DNA sequences at breakpoints were downloaded primarily from the UCSC genome browser. The Ensembl genome browser provided control comparison tests.

#### EBV genome comparisons

EBV binding locations on human chromosomes were obtained from publications [26-28], from databases, by interpolation from published figures, or by determining the location of genes within EBV binding sites. EBV binding data was based on lymphoblastoid and nasopharyngeal cancer cell lines. The NCBI BLAST program (megaBLAST) determined maximum homology scores between human DNA and all sequenced viruses in genome regions where EBNA1 binding occurred before anchoring EBV. When necessary, genome coordinates were all converted to the GrCH38 version. We further compared EBV binding sites to hereditary breast cancers with respect to chromosome break positions, epigenetic marks on chromatin, genes, and copy number variations. Locations of H3K9Me3 chromatin epigenetic modifications were from the MIT Integrated Genome Viewer (IGV) with ENCODE data loaded and from the UCSC genome browser. Data from the ENCODE website. (www.ENCODEproject.org) also provided positions of H3K9Me3 marks. Homology among viruses was determined by the method of Needleman and Wunsch [29].

#### Data analyses

Microsoft Excel, OriginPro, Visual basic, and Python scripts provided data analysis. Chromosome annotation software was from the NCBI Genome Decoration page and the Ritchie lab using the standard algorithm for spacing [30]. Statistical analyses used StatsDirect statistical software. Linear correlation, Kendall, and Spearman tests compared distributions of the same numbers of chromosome locations that matched viral DNA. Because the comparisons require the same numbers of sites, comparisons truncated data down to a minimum value of maximum homology (human DNA vs. viral DNA) of at least 400.

#### Genes associated with the immune response damaged in breast cancer

We compared breast cancer somatic mutations to the immune metagenome [31-34]. However, computerized lists of immune system gene functions vs. breast cancer mutations contain many potential uncertainties. Sets of genes involved in immune responses also mediate other functions and represent a vast and growing dataset (geneontology.org). Immune defense genes now extend to three different immune systems (intrinsic, innate, and adaptive). Sets of genes involved in cancer control by immune surveillance and immunoediting are not well characterized. To address these uncertainties, we determined whether mutated genes fit into known immune defense pathways. In addition, the Online Mendelian Inheritance in Man database (www.OMIM.org) was routinely consulted to determine gene function with frequent further support obtained through PubMed, Google scholar, GeneCards, and UniProtKB. The “interferome” was also sometimes used [www.interferome.org]. This extensive validation permitted including both direct and indirect effects of gene mutations.

## Results

Fig. 1 shows virus-human homology comparisons of one inter-chromosomal break-point in PD3945, a BRCA1 breast cancer (a) and one in a BRCA2 associated PD4874 a breast cancer (b). Panel (c) shows an intra chromosomal breakpoint. The comparisons include 450,000 bases extending in both directions from each breakpoint. The mega-BLAST program then compared the resulting 900,000 human base pairs around the breakpoint to all known viral sequences. Fig. 1 shows many DNA segments that are virtually identical to EBV variants (Human gamma-herpesvirus 4 variants, “HKNPC60” or “HKHD40”) at or near the different inter-chromosomal breakpoints shown. A chromosome fragment deleted in panel (C) includes virus-like sequences and has nearby viral matching sequences. Maximum homology scores between human DNA vs. herpes viral DNA are over 4000 for breast cancer PD3945 and just under 4000 for PD4874. Matches between human and viral sequences of about 97% for up to 2462 base pairs, with E “expect” values (essentially p-values) equal to 0. Many matches of shorter length in Fig.1 were also statistically significant.

**Fig.1.**
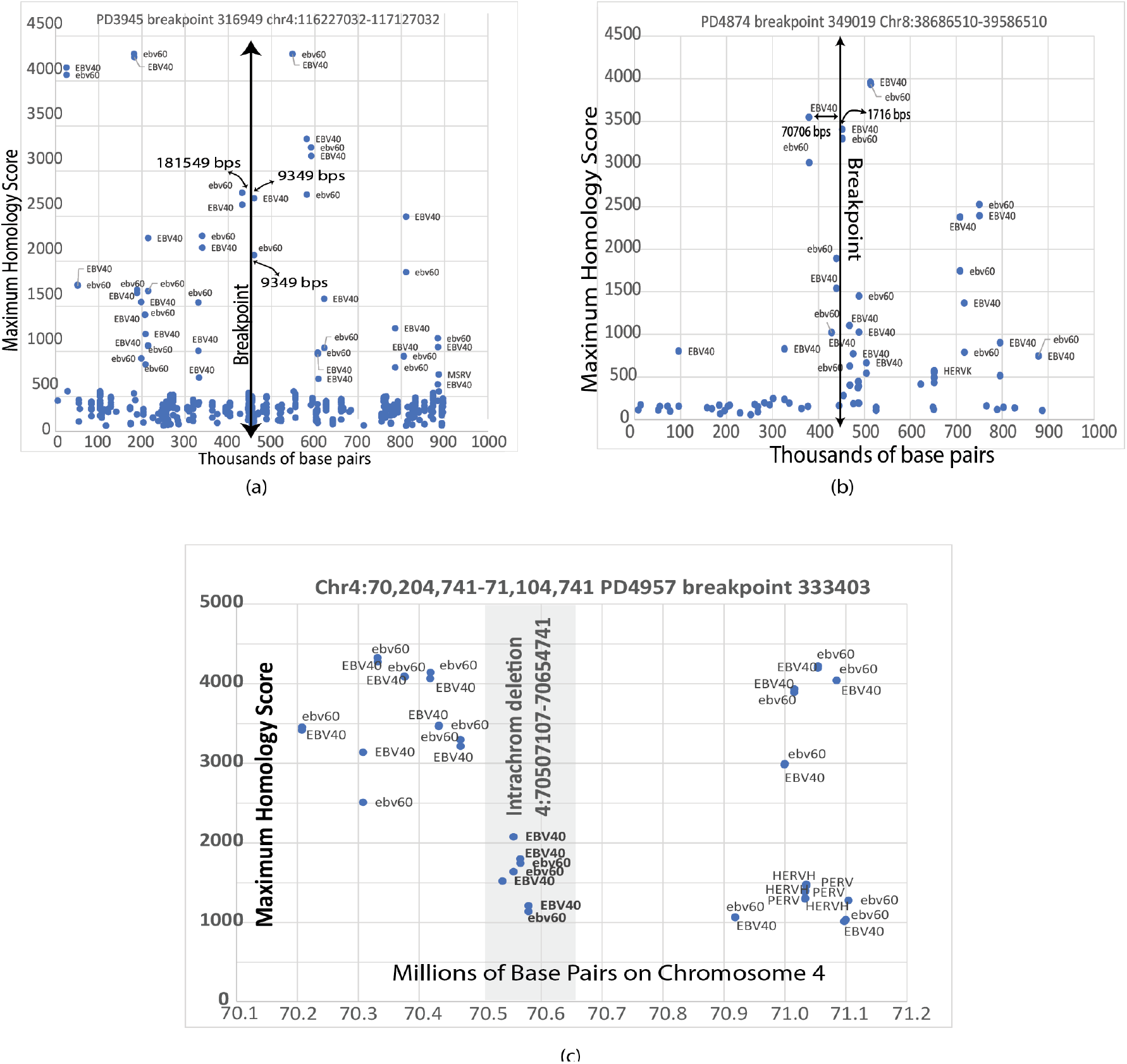
Examples of viral homologies at breast cancer breakpoints. Chromosome coordinates for locating the region with viral homologies are at the top of each panel. The lower horizontal axis represents distance from the breakpoint in thousands or millions of base pairs. The “y” coordinates are the maximum homology scores from megaBLAST analysis. The blue dots represent the start points of the homologous human-viral sequences. The numbers near breakpoints are the distances from the homology start points to the breakpoint. (a) A breakpoint on a BRCA1 associated breast cancer PD3945 is surrounded by DNA homologous to HKHD40 (EBV40) and HKNPC60 (ebv60). (b) A breakpoint in the BRCA2-associated breast cancer PD4874 is also near regions of homology to HKHD40 and HKNPC60. In the bottom panel (c), an intra-chromosome deletion on chromosome 4 (gray area) in BRCA2 associated breast cancer (PD4957) is near EBV40 and ebv 60 sequences and includes them. Weaker agreement with retroviral sequences is at greater distances at the lower right of the panel.

In Fig. 1, many human herpes viral sequences were within the viral DNA length (∼175,000 base pairs) of the linear lytic infection form. Bound latent EBV circular episomes could also interfere with the large complex apparatus required for complete DNA restoration by homologous recombination. Viral homologies suggest a large number of EBV binding sites with a substantial percentage at repetitive sequences. Lu et al. found 4785 EBV binding sites with over 50% overlapping a repetitive sequence element [26].

### Inter-chromosomal breakpoint distributions in BRCA2-Associated breast cancers

Fig. 2 shows the positions where inter-chromosomal breaks occur on 19 BRCA2 associated breast cancers, listed numerically at the bottom of the figure. There are significant differences in how the breakpoints distribute on different chromosomes. Spaces between breakpoints are roughly the same throughout chromosome 8. Breakpoints are much sparser on some other chromosomes, e. g., chromosomes 9, 12, and 21. Chromosome 8 was the site of the most numerous chromosome breaks. Inter-chromosomal breakpoints tend to cluster in specific chromosome regions for individual breast cancers. A few chromosome areas in Fig 2 show inter-chromosome rearrangements occur in only one cancer. Large numbers of base pairs often separate the breakpoint positions, indicated by horizontal lines on the chromosome diagrams. Breast cancer PD4956 has a large breakpoint cluster on chromosome 11, and PD4952 has a group on chromosome 14. PD8621 is one of only a few BRCA2-related breast cancers with breaks on chromosome 21.

**Fig. 2.**
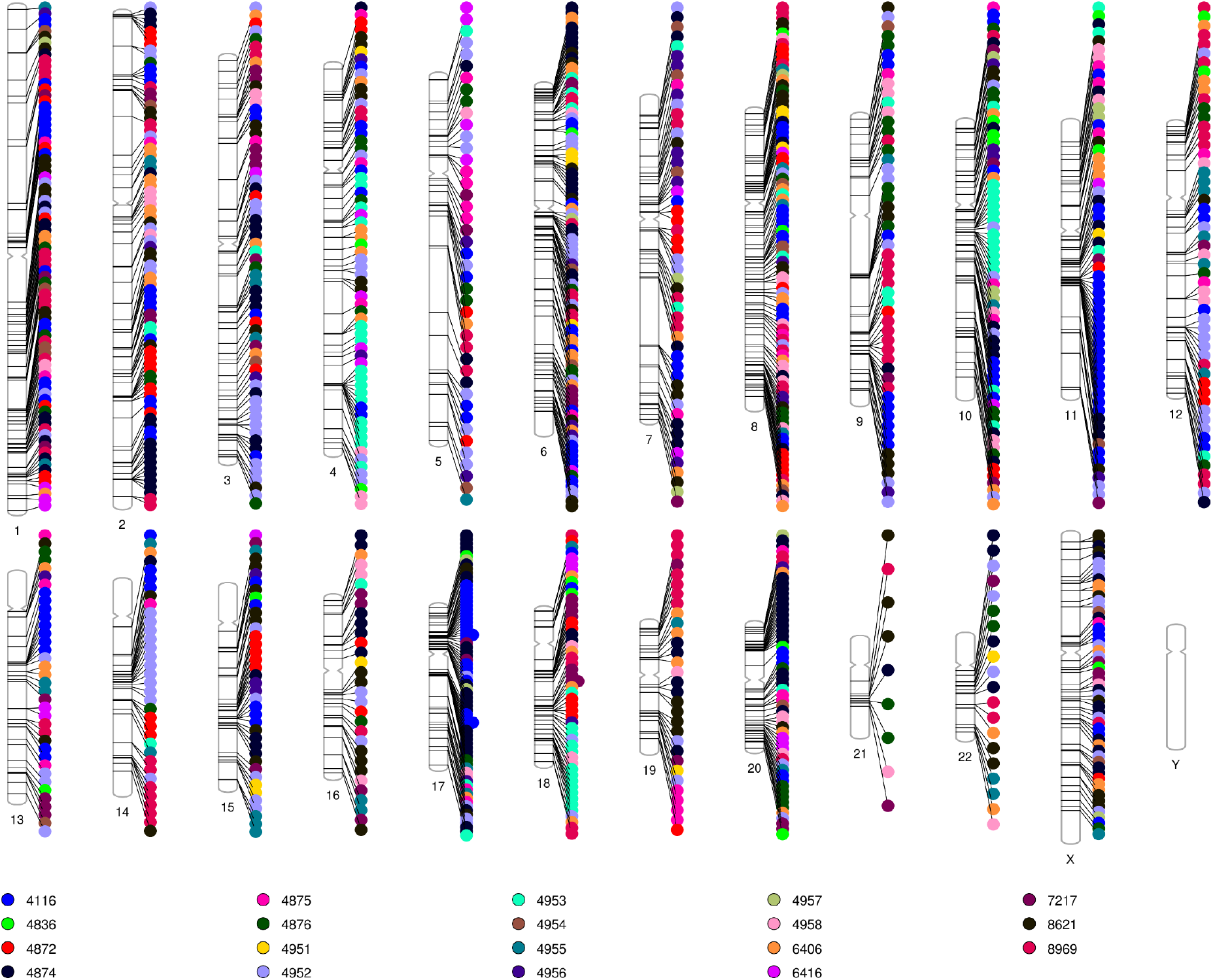
Distribution of chromosome breakpoints in BRCA2 mutation associated breast cancers over all human chromosomes. Each BRCA2 associated breast cancer (Table 1) is indicated by a four-digit number and labeled with a different color. Chromosome numbers are given below the chromosome drawing. The lines on the chromosome depictions indicate chromosome positions where the break occurs. The distributions are not random, with large regions of chromosomes that do not have breaks. Some breast cancer breaks appear to focus on a few areas and others are more widely distributed. Since all the breast cancers occurred in females, there are no breaks on the Y chromosome.

### Breast cancer breakpoint homologies to EBV are frequent

Chromosome 4 is one of the preferred landing sites for EBV infection. Breast cancer breakpoints cluster in inter-chromosome rearrangements on chromosome 4. To obtain an estimate of how often breakpoint regions have homology to herpes viruses, we tested every inter-chromosomal BRCA2 donor breakpoint on chromosome 4 for homology to herpes viruses. Nearly all (51/52) breakpoints had statistically significant homology to herpes viruses within about 200k base pairs (Fig. 3).

**Fig. 3.**
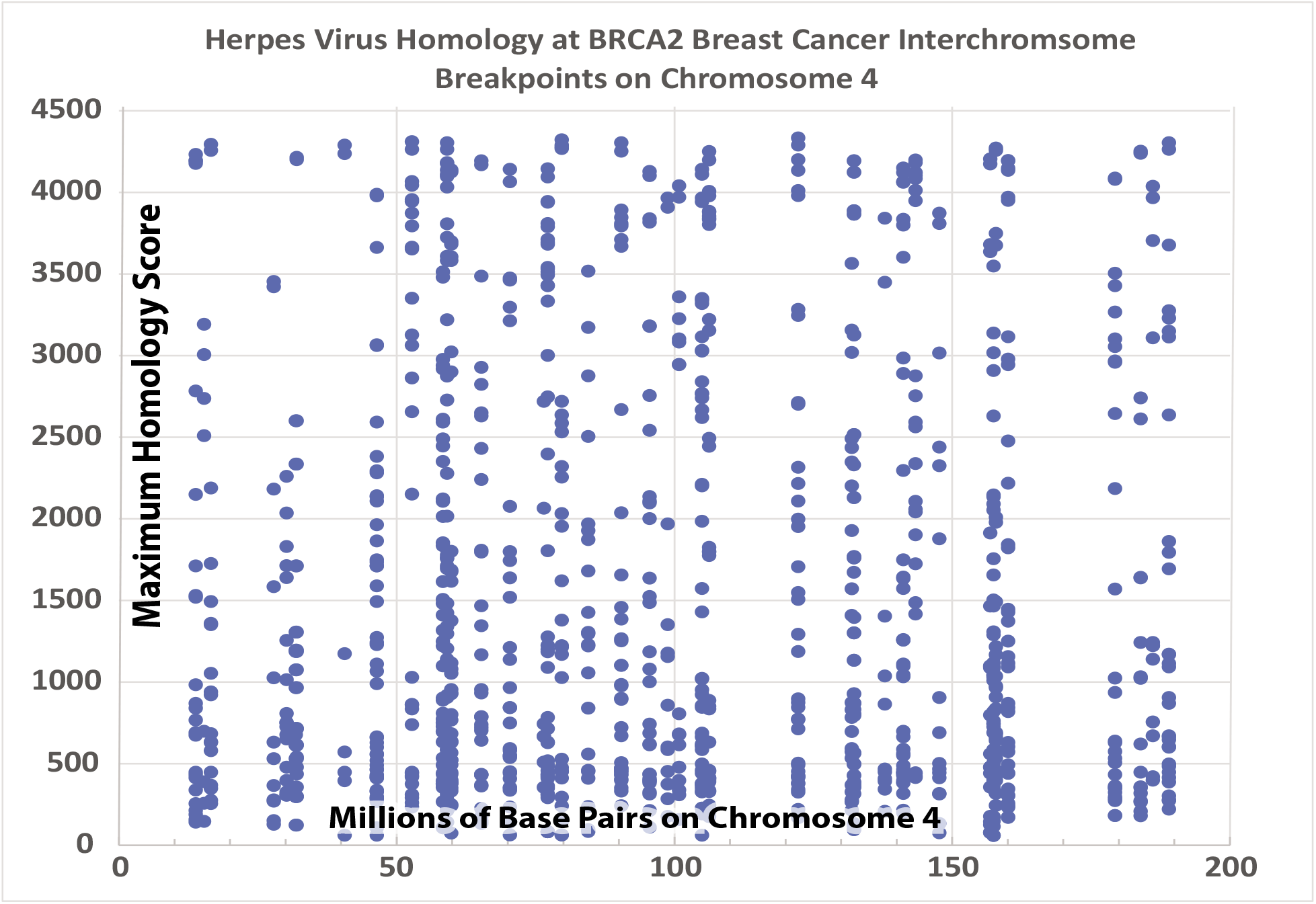
Statistically significant maximum homology scores (blue dots) for herpes virus DNA vs human DNA at inter-chromosome breakpoints on chromosome 4. Nearly all (51/52) inter-chromosome breakpoints on chromosome 4 in all BRCA2 associated breast cancers occur within 200K base pairs of EBV-like sequences. The value of 200K base pairs is approximately the length of the lytic EBV sequences allowing for some error and for tethering by EBNA1 dimers. The horizontal x axis represents the entire length of chromosome 4 and the Y axis shows the maximum homology scores. Values above 4000 represent about 97% homology over about 2500 base pairs. E (expect) values (essentially p values) = 0. Only homology scores over 100 are shown.

### Breast cancer breakpoint homologies to EBV are near to known EBV binding sites

#### Chromosome 2

Fig. 4 shows viral homologies in a 21 million base pair section of chromosome 2. The pink dotted lines in Fig. 4 indicate positions where EBV binding sites align with breast cancer breakpoints. We calculated R^2^ by comparing the coordinates of EBV binding sites to breakpoints. The R^2^ value of 0.91 estimates that EBV homology accounts for about 91% of the breakpoint position differences. The normal plot of the residuals from this analysis is approximately linear, suggesting that the residuals are distributed relatively randomly about zero.

**Fig. 4.**
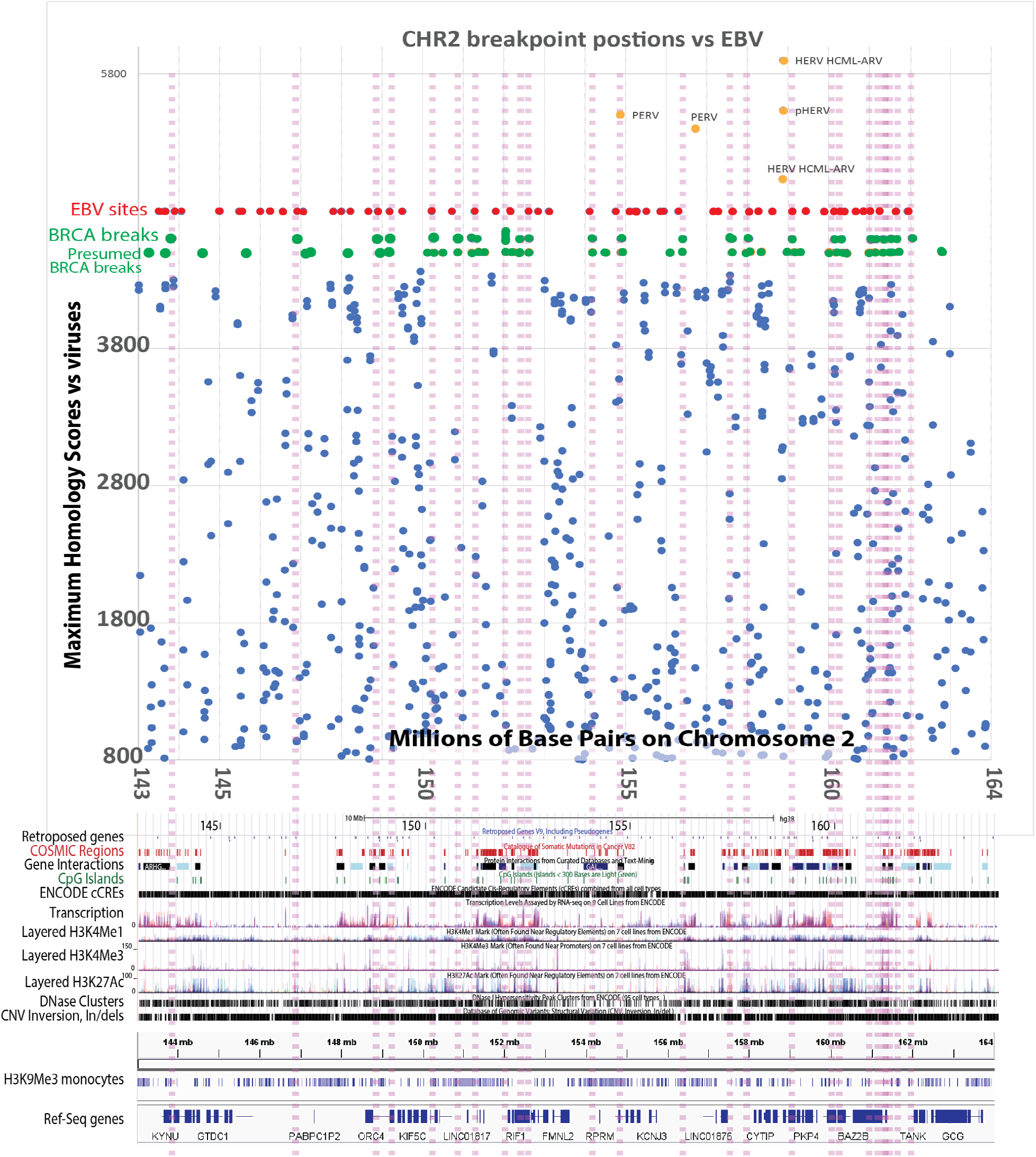
Breakpoints on 21 million bases in human chromosome 2 in BRCA1, BRCA2, and presumptive BRCA associated breast cancers (green dots) are near EBV anchors. Blue dots represent significant homology scores between human chromosome 2 base pairs vs. ebv60 and EBV40. Dashed lines align EBV anchor sites with BRCA and presumptive BRCA breakpoints. EBV anchor locations (red dots) and breast cancer breakages (green dots) are shown to allow comparisons. Porcine endogenous retrovirus and human endogenous retrovirus (PERV and HERV) variants (orange) may also contribute to breakpoints (top). The panel at the bottom shows that many of the breakpoints disrupt gene regulation, gene interaction, and transcription. Most breaks occur in regions that are accessible, according to DNase hypersensitivity. Most breast cancer breakpoints coincide with H3K9Me3 peaks in monocytes except for two break-points at about 153 million bps. Many breaks affect cancer-associated (COSMIC) genes, and many split reference genes within the region.

Human reference sequence genes in this region of chromosome 2 include KYNU, GTDC1, ACVR2A, KIF5C, STAM2, KCNJ3, ERMN, PKP4, BAZ2B, TANK, and DPP4 (bottom of figure). Chromosome breaks near these genes interrupt functions essential for immunity and preventing cancer. For example, KYNU mediates the response to IFN-gamma. TANK is necessary for NFKB activation in the innate immune system. DPP4 is essential for preventing viral entry into cells.

Most homologies are again to EBV variants, but porcine endogenous retrovirus (“PERV,” isolate PERV_VIRES_GX05_C1 env) [35] has two high scores. Three HERV sequences [36] including a pseudogene (pHERV), also have significant homology to human DNA. The porcine PERV sequence lies within a retroposed area on chromosome 2 close to EBV sequences, within 28 Kbps 5’ and 80 Kbps 3’ of EBV sequences. These EBV-like ends potentially generate homologies for retro-positioning and inserting the porcine sequences.

DNA surrounding many breakpoints had the EBV variant sequences (HKHD40 and HKNPC60) in both the donor and acceptor sequences. Even DNA sequences within intra-chromosomal deletions contained these variant EBV sequences (Fig. 1c)

#### Chromosome 12

On a 14 million base-pair section of chromosome 12, we aligned EBV binding sites on chromosome 12, breast cancer breakpoints, and homology between human sequences and tumor-related herpes viruses (Fig 5). All breakpoints are near regions of homology to the EBV variant tumor viruses HKNPC60 and HKHD40. Again, the maximum homology scores (over 4000) correspond to about 2500 bps that are 97% identical. All but one of the breakpoints is near an EBV anchoring site. A few homologies to human and porcine retroviruses are outside this region of chromosome 12. All the breakpoints appear to be accessible as indicated as DNase hypersensitivity.

**Fig.5.**
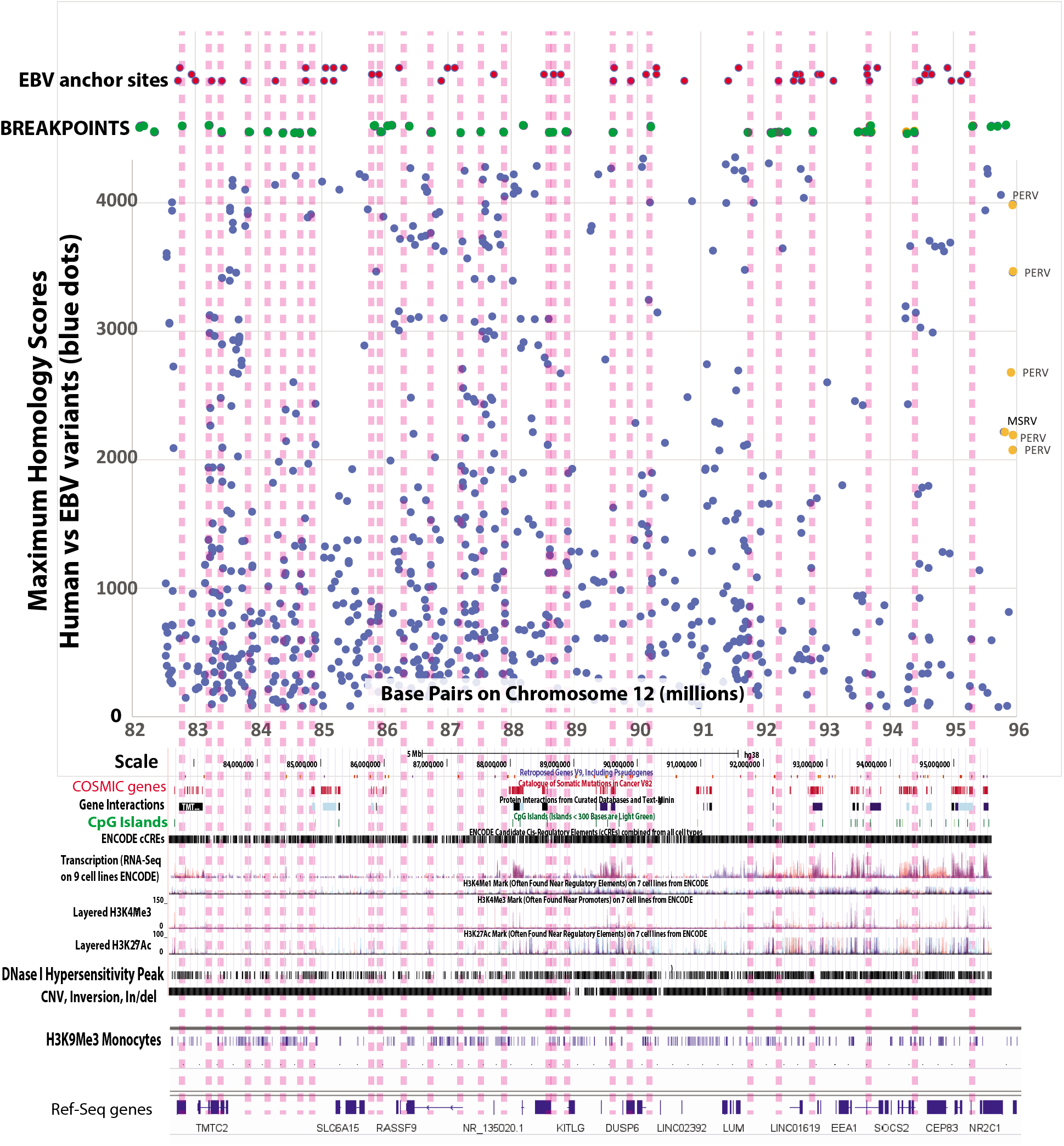
Breast cancer breakpoints on 14 million base pairs on chromosome 12 (green dots) in BRCA1, BRCA2, and presumptive BRCA associated (breast cancer at age <=35) occur in close proximity to EBV anchor sites. Statistically significant homologies to EBV viruses occur for all but one breakpoint within 200k base pairs of an anchor site. Green dots show the breast cancer breakpoints. Dashed vertical lines show that almost all breakpoints are near EBV tethering sites (red dots at top of the graph). The blue dots indicate positions of homology to the EBV variants HKHD40 and HKNPC60, but a few dots at the right match human and pork endogenous retroviruses. The bottom of the figure shows that breakpoints disrupt ENCODE candidate cis-regulatory elements (cCREs), H3K27Ac regulatory marks, and some regions of active transcription. Cancer-associated (COSMIC) genes and gene interactions are also disrupted. The breaks all occur in chromosome regions with accessible DNA (DNase hypersensitivity). Breakpoint positions also agree with positions of the EBV-associated epigenetic mark H3K9Me3 in CD14+ primary monocytes (RO-01946). The breaks usually disrupt one or more reference sequence genes, and the region gives rise to structural variation, i.e., CNV’s, inversions, and Insertion/deletions. Endogenous human and porcine retroviruses are less likely to affect the area and lie at the right.

All breakages disrupt regulatory properties shown in Fig 5. Results are consistent with data from epithelial cell lines showing EBNA1 binds cellular promoters to control cellular gene expression [37]. In the present work, the breakpoints all go through ENCODE candidate cis-regulatory elements (cCREs). Some breakpoints disrupt cancer-related (COSMIC) genes, transcription, gene interactions, and epigenetic marks such as H3K27Ac. At least 11 of the breakpoints disrupt reference genes (bottom of figure). The region also appears to be a focus for structural variation such as CNV’s, inversions, and short Insertion/deletions. The graph shows H3K9Me3 chromatin mark positions based on ChIP-Seq signals for monocytes. These markings are known to occur around EBV sites and support the positioning of EBV anchor sites. H3K9Me3 marks are thought to repress host transcription at EBNA1 binding sites [27], and the chromosome 12 region shown is rich in these sites. H3K9Me3 distributions on the host cell might also contribute significantly to EBV latency [27].

### Distributions of viral matches around different chromosome break-points correlate

We tested six different breast cancer breakpoints on different chromosomes to deterrent whether their separations from viral sequences had any correlation. The distribution of viral sequences from breakpoints along different chromosomes in various breast cancers is strongly correlated. The correlation held for linear regression (p=0.0093-0.000), Kendall (p<0.0001), and Spearman tests (p<0.0001).

### Identified EBV binding sites match breast cancer breakpoints

After converting the binding site coordinates to the same system as breast cancers, EBV binding sites [26] are close to early-onset breast cancer breakpoints and HKHD40 / HKNPC60 viral-human homologies (Fig 6). Different EBV binding sites on chromosome 6 and chromosome 1 (Including the CDC7 gene) were near an early-onset breast cancer break-point and HKHD40 / HKNPC60 viral-human homologies (Fig. 6). The known EBV binding site on chr5:141610249-141660249 was close to chromosome 5 breakpoints in breast cancer PD5945 (at 141,557,433 and 141,564,233). Intra-chromosomal breakpoints, which caused a deletion on chromosome 1, matched both the EBV variant sequences and the known anchor site. Fig. 6b shows the high-affinity anchor site for the EBV nuclear antigen (EBNA1) on chromosome 11[26]. The HDAC3 region on chromosome 5 did not contain nearby significant EBV variant homologies. Instead, there were lower maximum homology matches to retroviruses. The HDAC3 results suggest there are alternate mechanisms perhaps involving retroviral participation in breast cancer breakpoints. Nonetheless, viral binding sites were always near at least one breast cancer breakpoint. Five of the six localized EBV binding sites were near HKHD40 / HKNPC60 human viral homologies.

**Fig. 6.**
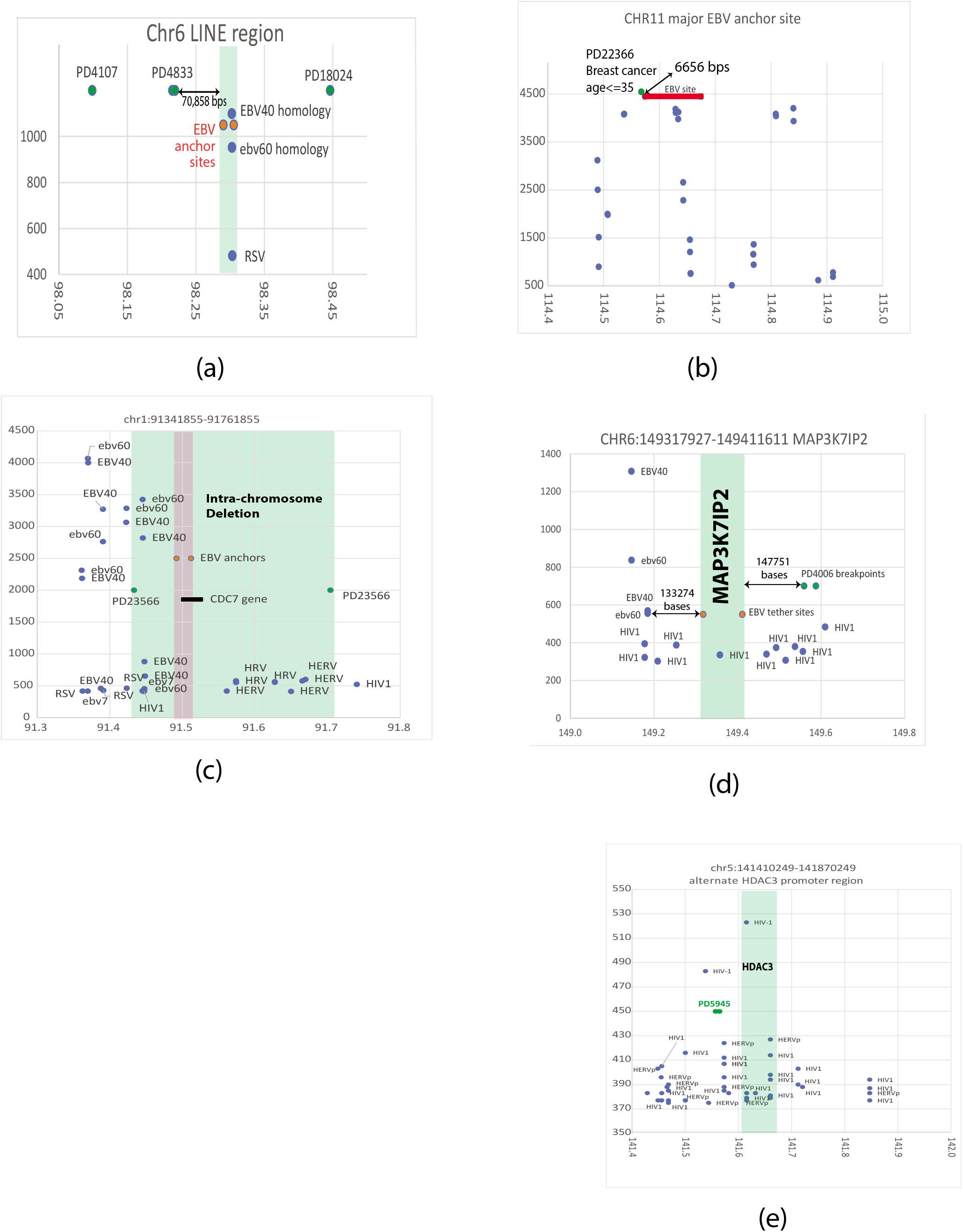
(a) A direct EBV binding site is near the region chr6: 98850000-98860000, which contains a LINE sequence (within the red dots) [26]. The area is near breakpoints in presumptive BRCA-associated breast cancers (green dots). Significant HKHD40 and HKNPC60 homologies overlap the LINE sequence. The break in breast cancer PD4833 is 70,858 bps from the EBV binding site. Weaker homology to the respiratory syncytial virus (RSV) also exists. In (b), a primary EBV binding site is within 6656 base pairs of a breakpoint in breast cancer PD22366. (c) EBV variant homologies near CDC7 fall within a deleted segment in an intra-chromosomal rearrangement on chromosome 1. Weaker retrovirus homologies are also shown. In (d), EBV anchors are within 148K bps of a breast cancer breakpoint. The MAP3K7IP2 gene has EBV anchor sites at its boundaries and nearby viral homologies at 133,274 bases. In (e), EBV binding near the HDAC3 gene does not have EBV variant homologies within 200K base pairs, but breakpoints in breast cancer PD5945 are still close to the region. Somewhat weaker homologies exist with HIV1 variants and endogenous retroviruses.

### Viral homologies around breakpoints in BRCA - associated breast cancers are difficult to distinguish from cancers caused by EBV

To compare breakpoints in BRCA1,2 associated breast cancers with model cancers known to be caused by EBV, we tested sequences around known recurrent gene fusions in nasopharyngeal cancer and Burkitt’s lymphoma (which has a 30-50% association with EBV). We compared the DNA within about 200k base pairs around breakpoints in these known EBV mediated cancers.

Fig. 7 shows gene fusions in nasopharyngeal cancer (as models for known EBV mediated cancers). These models resemble breast cancers in that they have homologies to EBV variants at breakpoints where fusions between genes occur. For example, the MAML2 gene, which rearranges in some NPC cancers, has homology to the EBV variants. The YAP1 breakpoint in its fusion to MAML2 occurs just within YAP1 gene boundaries. (Fig 7a). In Fig 7b, the PTK2 gene has strong similarities to EBV variants within its sequence. In Fig 7c, the RSRC1 breakpoint in the gene fusion RSRC1 to PTPLB is close to regions of strong EBV homology. In contrast, the breakpoint in PTPLB has weaker homology to endogenous and exogenous retroviral sequences. Five of the six gene fusions breakpoints are within the length of EBV variant number of base pairs from regions of EBV homology. In the PTPLB region of chromosome 3, there are numerous retroviral sequences near the breakpoint. Base pair distances from the EBV variant sequences are just outside the absolute values of the numbers of base pairs in the EBV variants.

**Fig. 7.**
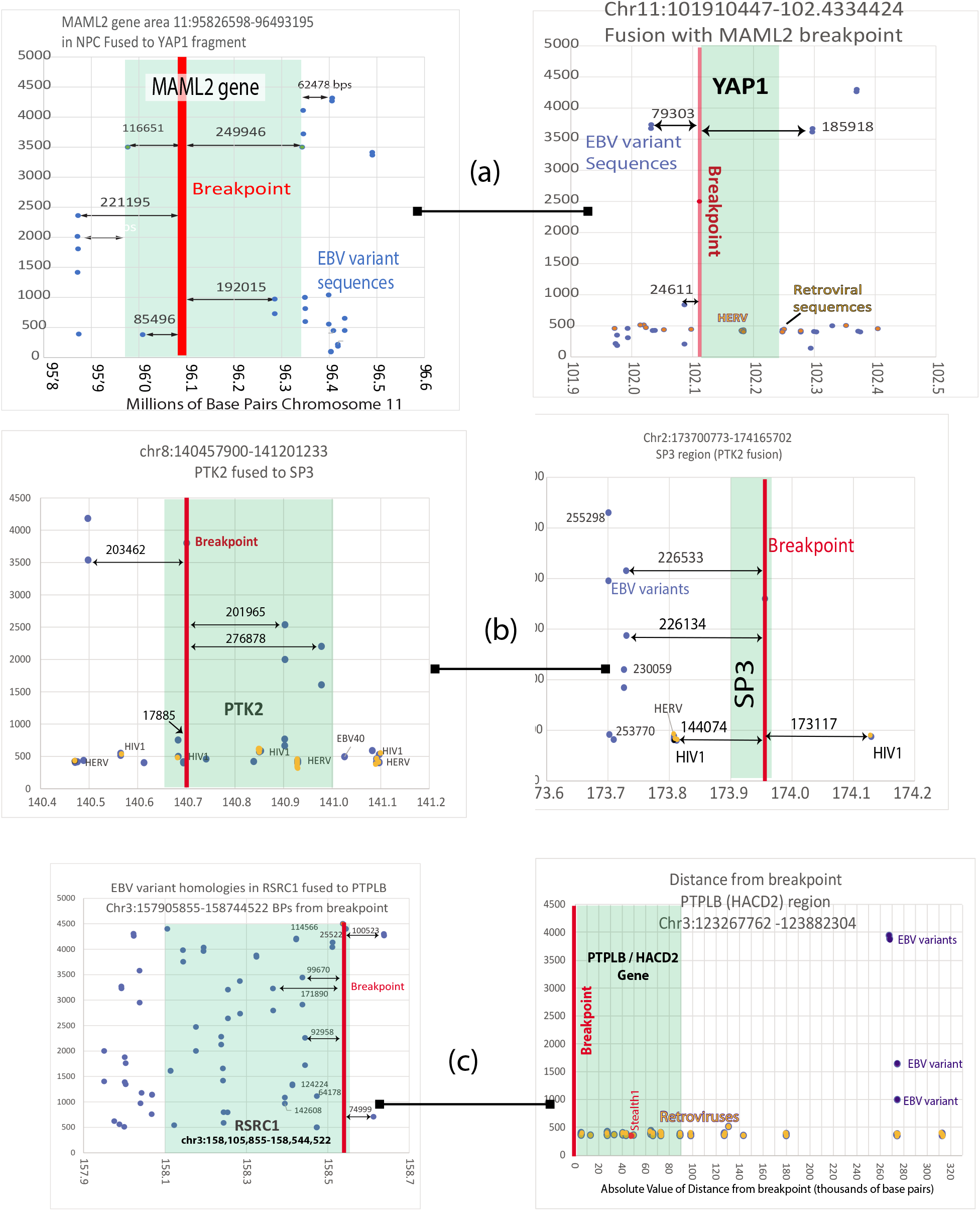
Gene fusions in nasopharyngeal cancers are near human sequences with strong homology to EBV tumor variants and sequences with weaker homology to retroviruses. Gene fusions are shown as left to right pairs (a) MAML2-YAP1, (b) PTK2-SP3, and (c) RSRC1-PTPLB). Maximum homology scores are plotted against chromosome locations for the first five panels. The PTLB panel (lower right) graphs maximum homology vs absolute valules for the total distance of viral homologies from the breakpoint. Orange dots indiate the start point of retroviral homologies.

Fig. 8 shows distances to EBV homologous regions starting from breakpoints in nasopharyngeal cancer NPC-5989 calculated from data from reference 12. In NPC-5989, all the breakpoints on chromosomes 1, 2, 8,and 11 are close to human genome regions that resemble the EBV variants (Fig. 8).

**Fig. 8.**
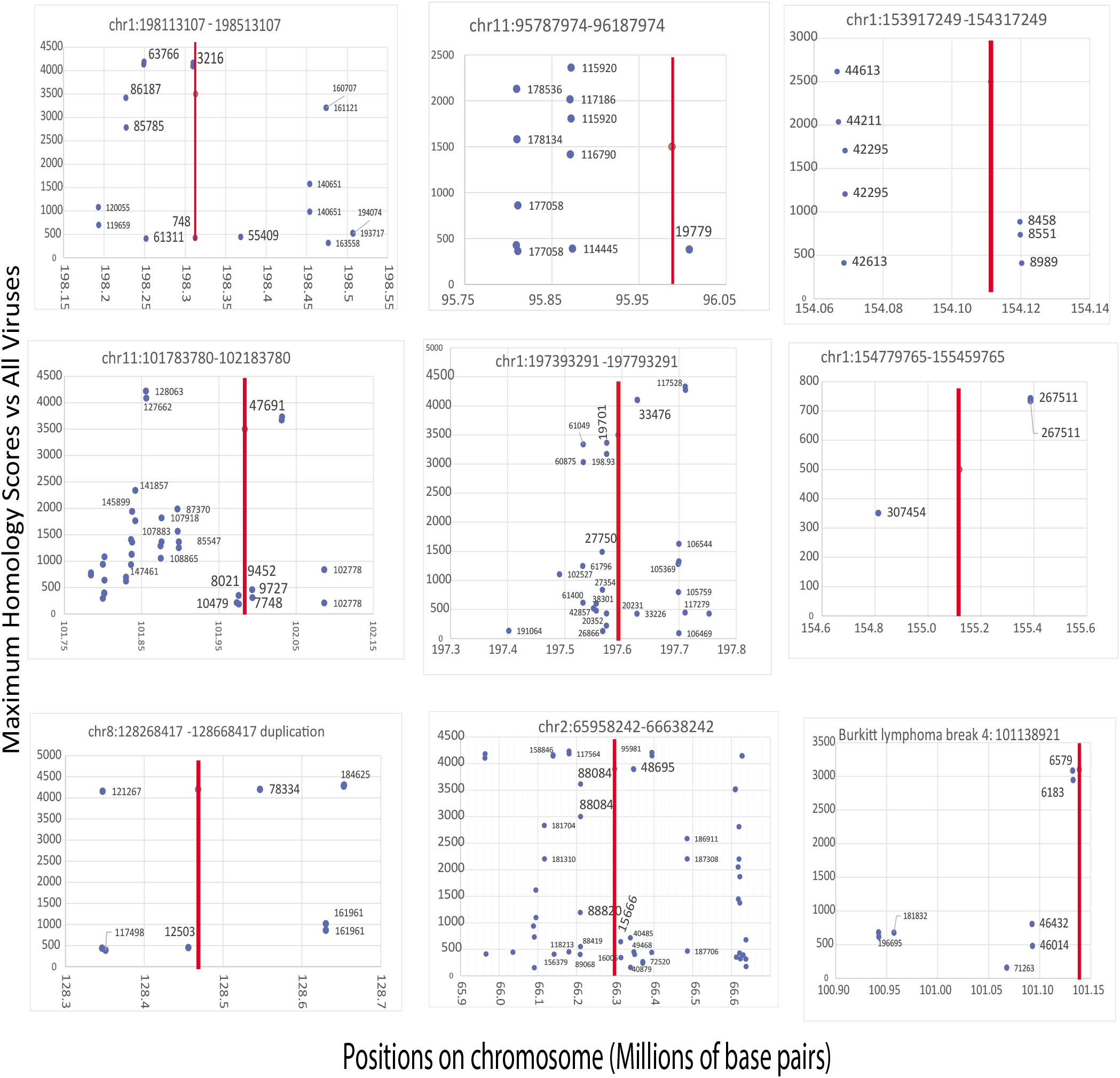
Distributions of EBV variant homologies around eight breakpoints (indicated by red lines) in nasopharyngeal cancer NPC-5989 [12] resemble distributions around hereditary breast cancers. Burkitt’s lymphoma break coordinates are also near DNA sequences with EBV variant homologies (lower right panel). The numbers near each of the blue dots are the number of base pairs between the breakpoint and the start of the homologous sequences. The numbers on the horizontal axis are the numbers of base pairs on the chromosome in millions. The vertical axes are again the maximum homology scores vs. all viruses

Table 2 shows that some breakpoints in NPC and BL cancers are near hereditary breast cancer breakpoints. Most (18/25) cancer breaks are within the number of bases in the EBV reference sequence (∼175,000). Comparable breakpoint positions in BRCA-breast vs. NPC vs. BL cancers differ by less than 1.25%). Table 2 also includes five values ranging from 181,246 to 274,186 base pairs. If the breakpoint’s actual value is closer to the average of the breast and other cancer breakpoints, then the break position still lies within the number of bases in EBV. Normality plots of the breakpoint positions for naso-pharyngeal cancer and Burkitt’s lymphoma vs. breast cancers are identical, go through zero, and follow the line y=x. One set of data can calculate the other using the equation Breast Cancers=1.00021NPC_BL +9012 (p<0.0001). An unpaired t-test showed differences between the two sets of data were not significant, p<0.001.

**Table 2.**
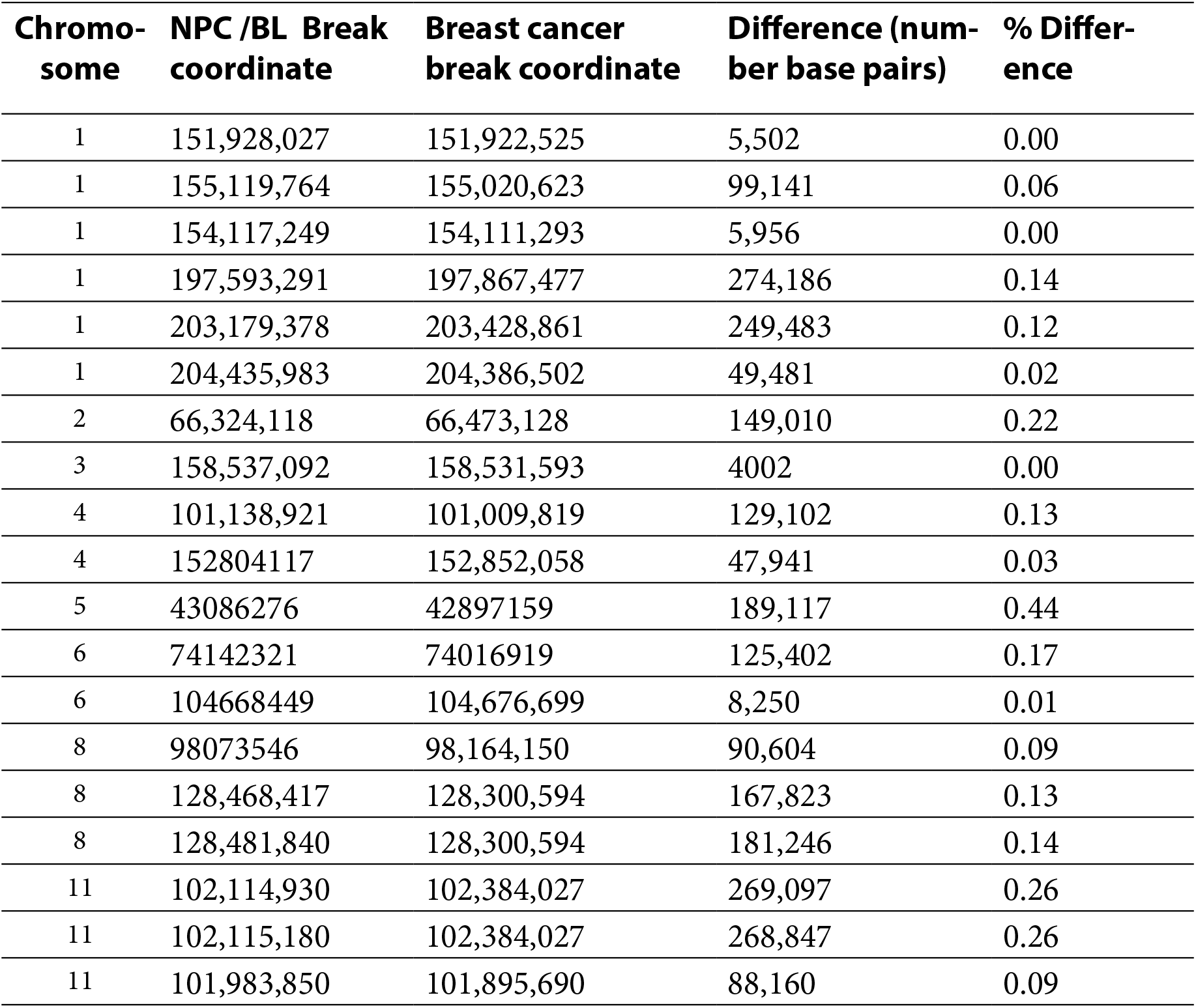
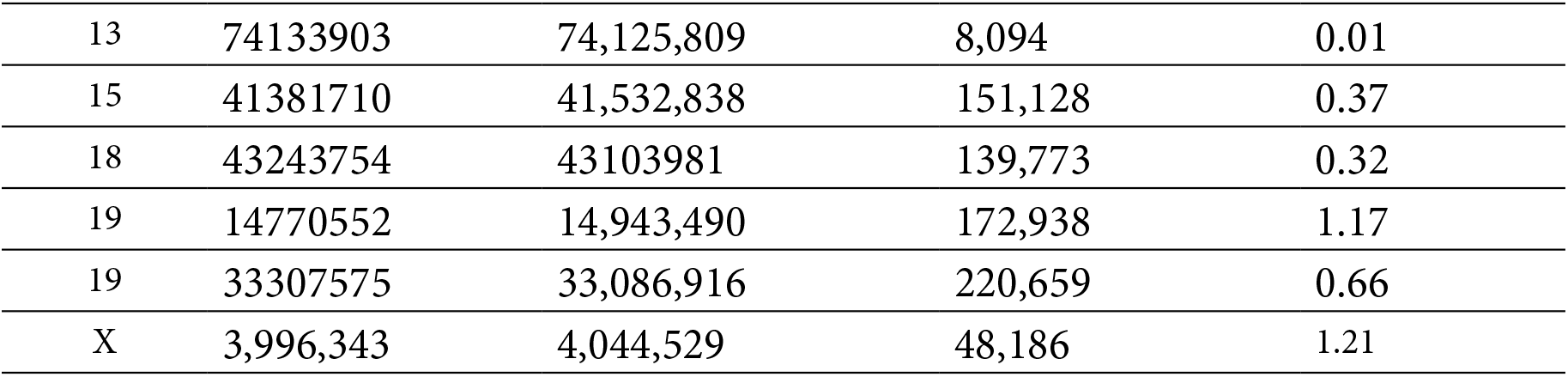
Comparisons of Cancer Chromosome Breakpoints.

### Damage to genes needed for the immune system in BRCA1 and BRCA2 associated breast cancers

All 25 of the hereditary breast cancers had significant damage to genes needed for immune system functions (Table 3). A total of 1307 immune-related genes had mutations. Table 3 shows the 20 most frequently mutated genes in the breast cancers belong to the “immune genome”. Of these top 20 genes, 8 (40%) belong to the innate immune system, and the remaining 12 participate in adaptive immunity. A variety of cells express the genes, including macrophages, B-cells, mast cells, helper- and memory-T-cells.

**Table 3.** The top 20 most commonly mutated genes in hereditary breast cancers are associated with the immune and nervous systems

| **Gene** | **Mutations in 25 BRCA1 or BRCA2 breast cancers** | **Type of Immunity in database [49]** | **Example of function in immunity** | **Known or likely connection to the nervous system** |
| --- | --- | --- | --- | --- |
| AKT3 | 20 | Innate | AKT3 amplifies innate immune responses to DNA or RNA virus infections [50]. AKT3 binds Interferon Response Factor 3, enhancing its activation and stimulating interferon responses. | ✓ |
| FRMD4A | 20 | Innate | Depends on Interferon Regulatory Factor 5 [51]. Affects antigen presentation and subsequent effects on dendritic cells. Mediated by cytoskeletal actin and endocytosis | ✓ |
| ARHGAP15 | 19 | Innate | Negative regulator of neutrophil function. Affects mast cell function | ✓ |
| CAMTA1 | 19 | Adaptive | Pattern of methylation distinguishes subsets of T-cells [52] | ✓ |
| DPYD | 19 | Innate | Interferon response [53]. Affects infection severity in mice. Natural killer cell function | ✓ |
| CHRM3 | 18 | Adaptive | Cholinergic receptor muscarinic 3 Deficiency associated with immune impairment [54] Targeting by miRNAs related to inflammation [55]. | ✓ |
| ADAMTS12 | 17 | Adaptive | Member of a group of related proteins with many connections to the immune system and immune disorders [56] | ✓ |
| DAB1 | 17 | Adaptive | Associated with microbial profile [57] | ✓ |

| Gene | Mutations in 25 BRCA1 or BRCA2 breast cancers | Type of Immunity in database [49] | Example of function in immunity | Known or likely connection to the nervous system |
| --- | --- | --- | --- | --- |
| ADAMTS3 | 16 | Innate | Member of a group of related proteins with many connections to the immune system and immune disorders [56] Mast cell function | ✓ |
| COL23A1 | 16 | Adaptive | Enriched in plasmacytoid dendritic cells | ✓ |
| DLC1 | 16 | Adaptive | Immune regulation of stem cells through interactions with NOTCH-1 [58] Type 2 T helper cell | ✓ |
| ADAM12 | 15 | Adaptive | Controls response to infectious disease [59]. Central memory CD8 t-cells | ✓ |
| CD36 | 15 | Adaptive | TLR4-TLR6-Cd36 activation is a common molecular mechanism by which atherogenic lipids and amyloid-beta stimulate sterile inflammation. Gamma delta T cell function | ✓ |
| CDH2 | 15 | Innate | Maintains stem cell quiescence [60]. Natural killer cell function | ✓ |
| DACH1 | 15 | Innate | Downregulated in lymphoid cell progenitor [61] | ✓ |
| CREB5 | 14 | Innate | CREB factors limit proinflammatory responses, provide anti-apoptotic signal, regulate Th1, Th2, Th3 responses, generate and maintain regulatory T cells [62] | ✓ |
| ANK1 | 13 | Adaptive | Type 17 T helper cell | ✓ |
| HDAC9 | 13 | Adaptive | Known to alter B-cell responses in autoimmune diseases Immature B cell function. | ✓ |
| COL4A1 | 12 | Adaptive | Central memory CD4 T cell | ✓ |
| F13A1 | 12 | Adaptive | Phagocytes at CNS borders [63] | ✓ |

Multiple components of immune defenses mutate in every one of 25 hereditary breast cancer genomes. Mutations also deregulate the immune response and contribute to cancer, e.g., by causing chronic inflammation. The mutations affect processes such as cytokine production, autophagy, etc. Every breast cancer had mutations affecting these functions or their regulation. The impaired functions depend on many genes dispersed throughout the genome, so mutation in only one gene can compromise a complex process. Each breast cancer genome has a different set of these mutations, with many genes only occasionally damaged in more than one breast cancer. Antigen and viral recognition, signaling essential to transmit the immune responses, and immune regulation can all be impaired. Damage affecting the nervous system was also universal. Some herpes viruses establish permanent infection within the central nervous system even after other sites clear [38].

Significant differences exist in distributions and numbers of mutations among the breast cancer genomes. Table 1 shows considerable differences in the numbers of DNA breaks [1,2,4,39]. Despite these differences, the significant common properties of many mutations are that they cripple some aspect of innate immunity, its regulation, or its connections to adaptive immunity. Damage to some genes can potentially deregulate immunity and inflammation in breast cancers (Table 3), damaging specific responses to antigens or pathogens. Deregulation could conceivably result from mutations alone without exposure to antigens or pathogens, impairing host response to breast cancer whatever its cause.

### Controls

#### Viral variants match other herpes viruses

Both HKHD40 and HKNPC60 viruses were typical of many other herpesvirus isolates, with some haplotypes conferring a high NPC risk [8]. About 100 other gamma herpes viral variants strongly matched both HKHD40 and HKNPC60 in regions with enough data to make comparisons possible. HKNPC60 was 99% identical to the EBV reference sequence at bases 1-7500 and 95% identical at bases 1200000-1405000. HKHD40 gave values of 99 and 98% identity for comparisons to the same regions.

If the infection is active and lytic, many breakpoints lie within the length of bases in the EBV genome and EBV anchor sites. Alternatively, viral episome binding may block homologous recombination machinery from precisely correcting DNA breaks if EBV infection remains latent.

#### Breast cancers have similarities to known or likely viral cancers

Positive controls found 1143 identical genes mutated in breast cancers vs. viral cancers of the cervix. About 50% of the genes mutated in a set of breast cancers were identical to genes mutated in the viral cancer Burkitt’s lymphoma and virally associated liver cancers. Human telomeric sequences did not have homology to EBV, e.g., DNA bases 1-1,000,000 on chromosome 11 had no homology to EBV.

The UCSC genome browser and the Ensembl genome browser gave indistinguishable results.

## Discussion

These results implicate variants of EBV in hereditary breast cancers because viral anchors match chromosome breakpoints; because many breast cancer chromosome breakpoints are near human sequences that match viruses closely; and because patterns of viral homology are difficult to distinguish from cancers known to be caused by EBV. Viral variants are significant breast cancer drivers without producing large numbers of active viral particles in the tumor. The association of EBV variant sequences with chromosome aberrations does not require the continuing presence of active virus anywhere within the resulting tumor. Because virtually everyone carries EBV infection, viral participation in breast cancer is very difficult to distinguish from its role in seemingly normal cells. Our comparisons of all virus sequences to the human genome reveal widespread, repeated EBV-like variant sequences dispersed throughout the human genome. In general, electron microscopy of EBV associated cancer cells does not detect EBV particles, but malignant cell nuclei do show viral DNA [40]

HKHD40 and HKNPC60 herpes viruses can exist as covalently closed circular episomes with multiple copies. These viral genomes assemble into chromatin, undergoing epigenetic histone and DNA modifications analogous to host genomes. This epigenetic programming of the viral episome controls the viral life cycle. The viruses switch from latent to lytic replication when signals from host cells indicate cell stress, DNA damage, another infection, or differentiation. In latently infected cells, the EBV protein EBNA1 binds host DNA and anchors viral episomes to host chromosomes at hundreds of sites.

Viral genomes remain attached to host chromosomes as parasites during mitotic cell division. This attachment maintains stable numbers of episome copies in multiplying host cells [41,42]. However, attached EBNA1 and viral particles also interfere with host chromosome replication. This interference becomes so significant in mouse models that B-cell tumors develop [43].

In infected B-cells, EBV establishes premalignant latent gene expression programs that also exist in lymphomas. On B-cell activation, the virus switches to its lytic replication program. During an early stage of EBV lytic replication, the virus generates strong nicks in host cell DNA, causing host chromosome aberrations [14]. EBV encodes a DNase (BGLF5) that causes these nicks and induces genomic instability in human epithelial cells, inducing chromosome micronuclei formation in epithelial cells [14]. BGLF5 also shuts off host protein synthesis and interferes with host cell DNA repair. The enzyme is a potent inducer of genomic instability. Lytic EBV replication was previously thought to destroy infected cells, which would prevent linear, complete viral DNA from causing tumors. However, recent work establishes that lytic, full-length EBV replication helps drive EBV-mediated malignancies [43]. Lytic replication also produces BALF3, another EBV nuclease and potential culprit [13].

The present study really began when we found EBV-like sequences very near many breakpoints in BRCA1 and BRCA2-associated breast cancer genomes. Fig. 1 shows three examples of breast cancer breakpoint positions close to human genome sequences that strongly resemble viral variants. These viral similarities also exist within intra-chromosome regions that get deleted in individual breast cancers (Fig.1 bottom). BRCA1 and BRCA2 mutation carriers are likely to be highly susceptible to EBV variant nucleases, interference with replication, and recombinase activity. Mutations in BRCA1 and BRCA2 genes inactivate the high-fidelity repair of EBV-induced chromosome damage.

Two explanations must be explored for the role of EBV-like variants in c;hromosome breaks in hereditary breast cancers. (1) Chromosome breaks in hereditary cancers occur because human DNA homologies to EBV-variants help guide viral variant binding in the epithelium of breast ducts, where viral nucleases damage the host chromosome. Alternatively, (2) pre-exising viral variant binding interferes with access to repair machinery so that breaks, whatever the cause, are slow to repair. In either event BRCA1 and BRCA2 gene deficits make repairs less accurate.

Of course, there are countless unrelated agents and mechanisms known to break DNA and chromosomes that do not involve viruses. Accordingly, chromosome breaks could easily follow some random distribution. In contrast, inter-chromosome BRCA1 and BRCA2 breakpoints are not distributed randomly among chromosomes, and there are marked differences among breast cancers in the positions where these breakpoints occur (Fig. 2). These different distributions suggest that inter-chromosome breaks followed by incorrect repairs do not result from random processes but have some selectivity.

A consequence of this selectivity is that nearly all the inter-chromosome break-points on the entire length of chromosome 4 occur within the approximate number of base pairs in lytic EBV variant sequences (roughly 200k in Fig. 3). Moreover, many breast cancer breakpoints are separated from viral homologous sequences by numbers of base pairs consistent with circular viral episomes (roughly 65000). Circular viral episomes could also be a dominant carcinogen.

Comparisons to actual sites of EBV anchoring by EBNA1 further supports a relationship between EBV and chromosome breaks in hereditary cancers. Comparisons on chromosome 2 (21 million base pairs) and chromosome 12 (24 million base pairs) give similar results (Figs 4-5). Short DNA comparisons for a few known EBV binding sites also show that viral binding, breast cancer breaks, and viral human homologies are near each other (Fig. 6). EBV anchors and EBV variant homologies fall within a deleted segment of chromosome 1 (shaded area) in early-onset breast cancer PD23566. Breakpoints on chromosome 6 from three breast cancers (PD4107, PD4833, and PD18024) surround EBV anchor sites and EBV variant homologies. A primary binding site on chromosome 11 [26] matches a breakpoint in an early-onset breast cancer (Fig. 6). Weaker EBV binding sites on chromosome 11 also agree with chromosome break positions in BRCA1 and BRCA2 breast cancers (data not shown).

Figs. 4-6 are consistent with EBV variants interfering with transcription. Chromosome sequences near EBV anchor sites are enriched in transcribed DNA, DNase accessible sequences, and regulatory sequences. EBNA1 has linking modules that form aggregates, but the linkers also bind EBV episomes to metaphase chromosomes. In many cases, the positions of EBNA1 anchors, breast cancer breakpoints, and human regions of EBV homology all agree.

EBV binds to human chromosomes using EBNA1-dimers as anchors for subsequent viral attachment [44,45]. The dimerized anchor itself can alter human chromosome structure and mediate distant interactions related to transcription during infection. EBNA1 anchor sites favor some repetitive elements, especially LINE 1 retrotransposons (Fig. 6), and correlate weakly with histone modifications [26]. EBNA1 binds close to transcription start sites for cellular genes and positively regulates them. These genes include MAP3K71, CDC7, and HDAC3. Fig. 6 shows that most of the EBV anchor sites near these genes are relatively close to chromosome breaks in breast cancers. HDAC3 on chromosome 5 is an exception. A region of chromosome 5 containing an EBV anchor site near HDAC3 does not have significant EBV variant homologies near an EBV binding site. Instead, there are much shorter and weaker nearby similarities to retroviruses.

Chromosome 12 shows strong homologies to porcine endogenous retroviruses just outside the test region. Data from chromosome 5 and chromosome 12 suggest that retroviruses may also participate in breast cancer breaks in agreement with much previous literature. Thoroughly cooking pork products should avoid danger from porcine endogenous retroviruses. However, despite assertions that xenotransplantation with pig cells is safe, as many as 6500 bps in human chromosome 11 are virtually identical to pig DNA. This extensive homology challenges the safety of transplantation from porcine donors.

Nasopharyngeal cancer (NPC) and Burkitt’s lymphoma (BL) have known links to EBV. At least 25 breakpoints in these known EBV mediated cancers are probably within experimental error [46] of breakpoint positions in breast cancers (Table 2). Running viral human homology comparisons for such accepted EBV-related cancers gave results that were difficult to distinguish from those obtained for breast cancers (Figs. 7-8). Five of six genes involved in gene fusions are near EBV variant sequences and include EBV-like variant sequences within the gene boundaries. The PTPLB gene has these EBV variant homologies at about 265-275,000 base pair distances with short, weak retroviral homologies much closer to the fusion breakpoint (Fig 7).

Breast cancer mutations cripple the immune system’s ability to control cancer-causing infections, remove cells damaged by disease, and regulate inflammation. Many genes mutated in human breast cancers link to some variable function within the immune system or structural barrier defenses [23,47,48]. Many mutations independently link to various infections, including all known cancer-causing microorganisms. The top 20 mutated genes in BRCA1 and BRCA2 breast cancers are all associated with some function of immunity (Table 3).

Mutational damage to genes essential for an immune response and structural barriers to tumor viruses facilitates infection. The relationships of the top 20 mutated genes to the nervous system may also favor herpes viral infections. Damage to the immune system in hereditary cancers suggests DNA breakages might result when viruses escape from control. For example, escape of EBV from latency would release nucleases that fragment human DNA. Moreover, abundant viral DNA might form heteroduplexes with pieces of human DNA that resemble viral sequences. EBNA1 anchor sites might facilitate interactions that lead to chromosomal translocations.

The finding of a relationship between breast cancer chromosome breaks and viruses is potentially actionable. Some immunotherapy strategies rely on augmenting the immune response, but this approach may need modification because mutations create additional holes in the immune response. The present evidence adds support for developing EBV treatment and a childhood herpes vaccine. EBV causes about 200,000 cancers per year of multiple different types. The prospects for producing an EBV vaccine are promising, but the most appropriate targets are still controversial.

## Notes

### Competing Interest Statement

The authors have declared no competing interest.

